# Fat body-specific reduction of CTPS alleviates HFD-induced obesity

**DOI:** 10.1101/2022.05.12.491743

**Authors:** Jingnan Liu, Yuanbing Zhang, Youfang Zhou, Qiao-Qi Wang, Ji-Long Liu

## Abstract

High fat diet (HFD)-induced obesity is a multi-factorial disease including genetic, physiological, behavioral, and environmental components. *Drosophila* has emerged as a useful model of metabolic diseases. Cytidine 5′-triphosphate synthase (CTPS) is a key enzyme for the *de novo* synthesis of CTP, governing cellular level of CTP and phospholipid synthesis. We have demonstrated that CTPS show the capacity to assemble into large filaments termed cytoophidia, which are evolutionarily conserved in bacteria, archaea and eukaryotes. Here, we show that CTPS acts in fat body to regulate body weight and starvation resistant in *Drosophila*. HFD-induced obesity elevates CTPS transcription and elongates cytoophidia in larval adipocytes. Fat body-specific depletion of CTPS alleviated HFD-induced obesity including body weight gain, lipid storage and TAG level. Moreover, a dominant negative form of CTPS reduces lipid accumulation and down regulates lipogenic genes. Therefore, our data not only provide a functional link between CTPS and lipid homeostasis, but also highlight the potential application of manipulating CTPS in the treatment of HFD-induced obesity.

## Introduction

Obesity has been the world-wide epidemic disease for decades. Obesity is defined as abnormal or excessive fat accumulation, which is a threat to health. Clinically, High fat diet (HFD)-induced obesity is a major risk factor for chronic diseases, including diabetes, cardiovascular diseases and cancer, which contributes to approximately 2.5 million deaths annually[1]. It has been shown that obesity caused by excessive dietary fat consumption and the harmful effects of its genetic factors are very important for understanding the mechanism of obesity and its related secondary diseases (such as nonalcoholic fatty liver) [2]. However, the interactions of genetic predisposition with environmental and lifestyle factors in the etiology of obesity remain to be fully elucidated.

CTPS is a key rate-limiting enzyme for *de novo* synthesis of the CTP metabolic pathway, catalyzing the ATP dependent transfer of the amide nitrogen from glutamine to the C-4 position of UTP to form CTP [3, 4]. In 2010, we and other groups reported that CTPS polymerizes into filamentous structures, termed as cytoophidia [5] or CTPS filaments [6, 7], in fruit flies [5], bacteria [6] and yeast cells [7]. Subsequent studies confirmed that the cytoophidium also exists in human cells [8, 9], plants and archaea [10–13], demonstrating that filamentation of CTPS is highly conserved across prokaryotes and eukaryotes. Multiple studies show that diversiform functions of CTPS cytoophidia in modulating its enzymatic activity [14, 15], maintaining cell morphology [6], and stabilizing CTPS protein [12, 16]. CTPS cytoophydia were found in various human cancers including hepatocellular carcinoma [17]. Although eventually a role for CTPS cytoophidia as a causative factor in these diseases has yet to be fully elucidated but it is thought to contribute to tissue homeostasis by coordinating cell growth/proliferation and nutrient availability. Notably, a large amount of CTPS is produced in mammalian adipose or hepatic tissues. However, the physiological function of CTPS in lipid homeostasis remains an open question.

*Drosophila* has emerged as a powerful and simplified model of metabolic diseases including HFD-induced obesity, diabetes and heart disease [18–21]. It is attractive for determining links between genetics, diet, and metabolism. *Drosophila* fat body, an organ with great biosynthetic and metabolic activity and conserved signaling pathways, plays an essential role in sensing nutritional conditions and responding with the integration of the lipid metabolism, which acts as a counterpart of mammalian liver or adipose tissue [22, 23]. Importantly, the single ortholog of CTPS in *Drosophila*, appears highly conserved, forming cytoophidia in larval adipocytes [24].

Employing *Drosophila* model, here we have uncovered physiological function in lipid metabolism and metabolic adaptation of CTPS cytoophidia response to HFD exposure. We find that HFD feeding induces elevated CTPS transcription and elongates CTPS cytoophidia in larval adipocytes. We provide *in vivo* evidence showing that depleting CTPS prevents weight gain and restricts adiposity. Our findings suggest that CTPS is harnessed by adipocytes to modulate lipid metabolism for metabolic adaptation and disease.

## Results

### Fat body-specific knowdown of CTPS leads to body weight loss

To examine the potential role of CTPS in fat deposition and obesity, we initially compared the body weight of flies harboring with CTPS deficiency in different tissues to wild type ones. We knocked down the expression of *CTPS* globally by *Tublin GAL4* driver and found that the body weight of the female with *Tub G4>CTPS-Ri* (1.083 mg; S.E.M.: ± 0.015 mg) was 8.45% less than that of the *Tub G4>+* (1.183 mg; S.E.M.: ± 0.043 mg) (Figure 1A). The body weight of the male with *Tub G4>CTPS-Ri* (0.775 mg; S.E.M.: ± 0.020 mg) was 6.06% less than that of the *Tub G4>+* (0.825 mg; S.E.M.: ±0.010 mg) (Figure 1A).

**Figure 1.**
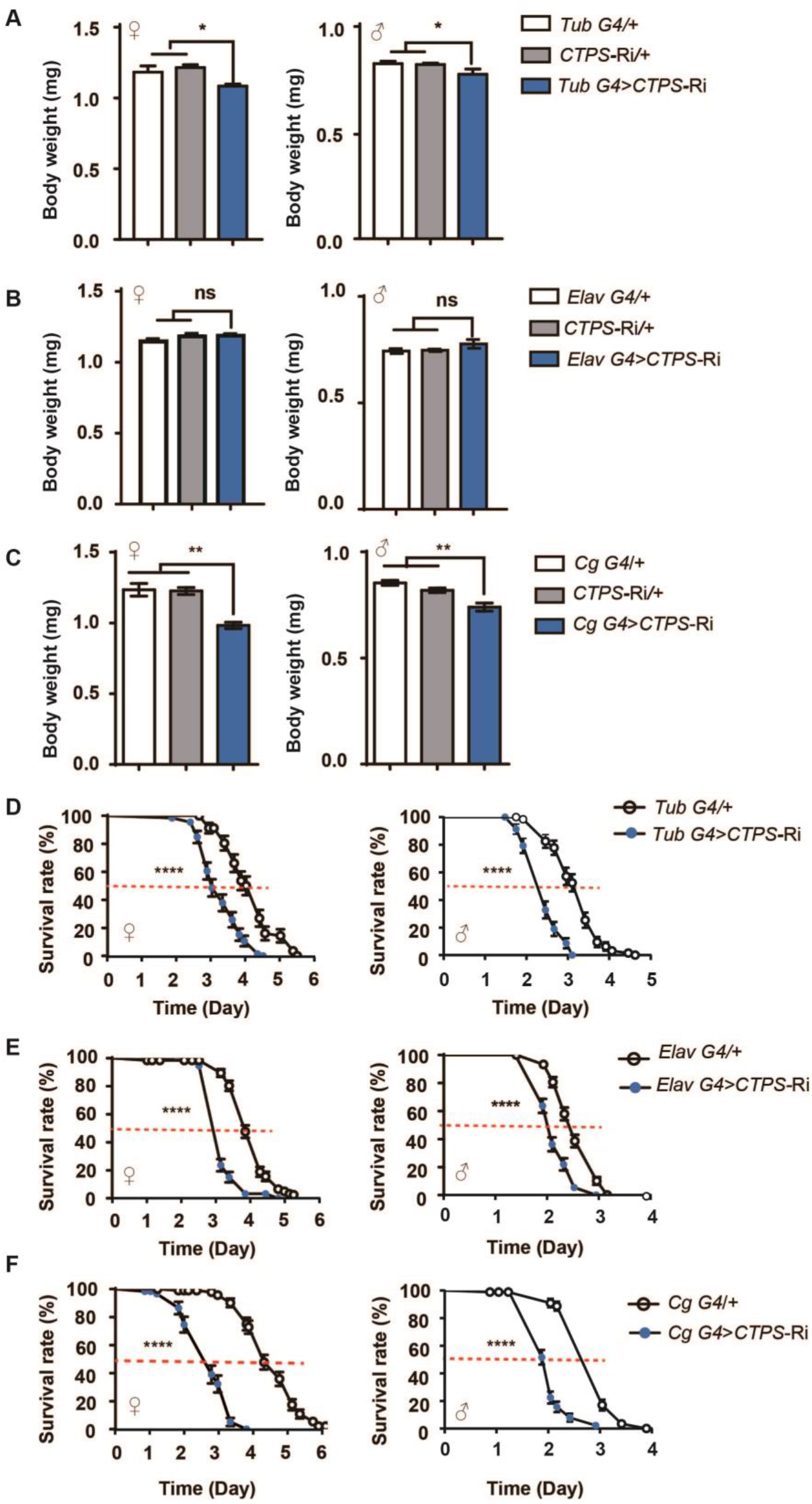
CTPS knockdown in fat body leads to body weight loss. (**A**-**C**) Body weight of 5-day old adults fly from the indicated genotypes. (30 flies/group, 5-6 groups/genotype, 3 biological replicates). *Tub G4>CTPS* Ri versus *Tub G4>* + or *CTPS*-Ri /+ (A), *Elav G4>CTPS-Ri* versus *Elav G4>* + or *CTPS*-Ri /+ (B), *Cg G4>CTPS-Ri* versus *Cg G4>* + or *CTPS*-Ri/+ (C). All values are the means ±S.E.M. ns, no significance, **\*** p < 0.05, **\*\*** p < 0.01, by one-way ANOVA with a Tukey *post hoc* test. (**D**-**F**) Survival rates are measured for starved female and male adult flies from the indicated genotypes (5 days of age; 30 flies/group, 5 groups/genotype, 3 biological replicates). *χ*^2^=35.6 for female, *χ*^2^=61.0 for male, *Tub G4>+* versus *Tub G4>CTPS-Ri*, ****p < 0.0001 by log-rank test (D), *χ*^2^=97.8 for female, *χ*^2^=54.4 for male, *Elav G4>+* versus *Elav G4>CTPS-Ri*, ****p < 0.0001 by log-rank test (E), *χ*^2^=152 for female, *χ*^2^=104 for male, *Cg G4>* + versus *Cg G4>CTPS-Ri*, ****p < 0.0001 by log-rank test (F).

Then, we knocked down the expression of CTPS in central neuron system by *Elav GAL4*, the body weight of both female and male with *Elav G4>CTPS-Ri* was no obviously different compared with that of the *Elav G4*>+control lines (Figure 1B).

Importantly, when specific knockdown of the expression of *CTPS* by *Cg G4*, a fat body GAL4 driver, the body weight of the female with *Cg G4>CTPS-Ri* (0.983mg; S.E.M.: 0.031±mg) was significantly reduced and less than 20.4% that of the *Cg G4>+* (1.235mg; S.E.M.: 0.062±mg) control line (Figure 1C). Similarly, the body weight of the male with *Cg G4>CTPS-Ri* (0.740mg; S.E.M.: ±0.027mg) was also significantly reduced and 13.2% less than that of the *Cg G4>+* control line (0.853mg; S.E.M.: 0.016±mg) (Figure 1C).

### CTPS is required for starvation resistence

An increase in body weight means an increase in energy reserves particularly in lipid stores, which seems to be a common adaptation to starvation in laboratory experiment [25]. We then asked whether CTPS affects the starvation response. We found that both female and male *Tub G4>CTPS-Ri* flies lived much shorter and showed 32.6% and 28.1% decreases in their median survival rates, respectively (Figure 1D). The *Elav G4>CTPS-Ri* fly also exhibit dramatic defects in survival during food deprivation, female and male show 28.1% and 15% induction in their median survival rates, respectively (Figure 1E). Notably, both female and male *Cg G4>CTPS-Ri* flies exhibited shortening of survival, 33.3% and 32.1% decreases in their median survival rates, respectively (Figure 1F).

### CTPS in the fat body are crutial for body weight maintenance

Considering potential leaking expression, we also knocked down *CTPS* expression using another fat body driver line, *PPL G4*. Consistently, the line showed similar decreases of body weight and survival during food deprivation (Figure S1A and B).

We further investigate the body weight and fat mass in larvae. Similarly, body weight of larvae was dramatically reduced in *Cg G4>CTPS-Ri* versus *Cg G4>+* lines (Figure 2A). Floating rate reflects larval fat mass [20]. We found that 90% larvae of *Cg G4>CTPS-Ri* sank down to the bottom of vial while 80% of *Cg G4>+* larvae floating on the top of 9% sugar solution (Figure 2B).

**Figure 2.**
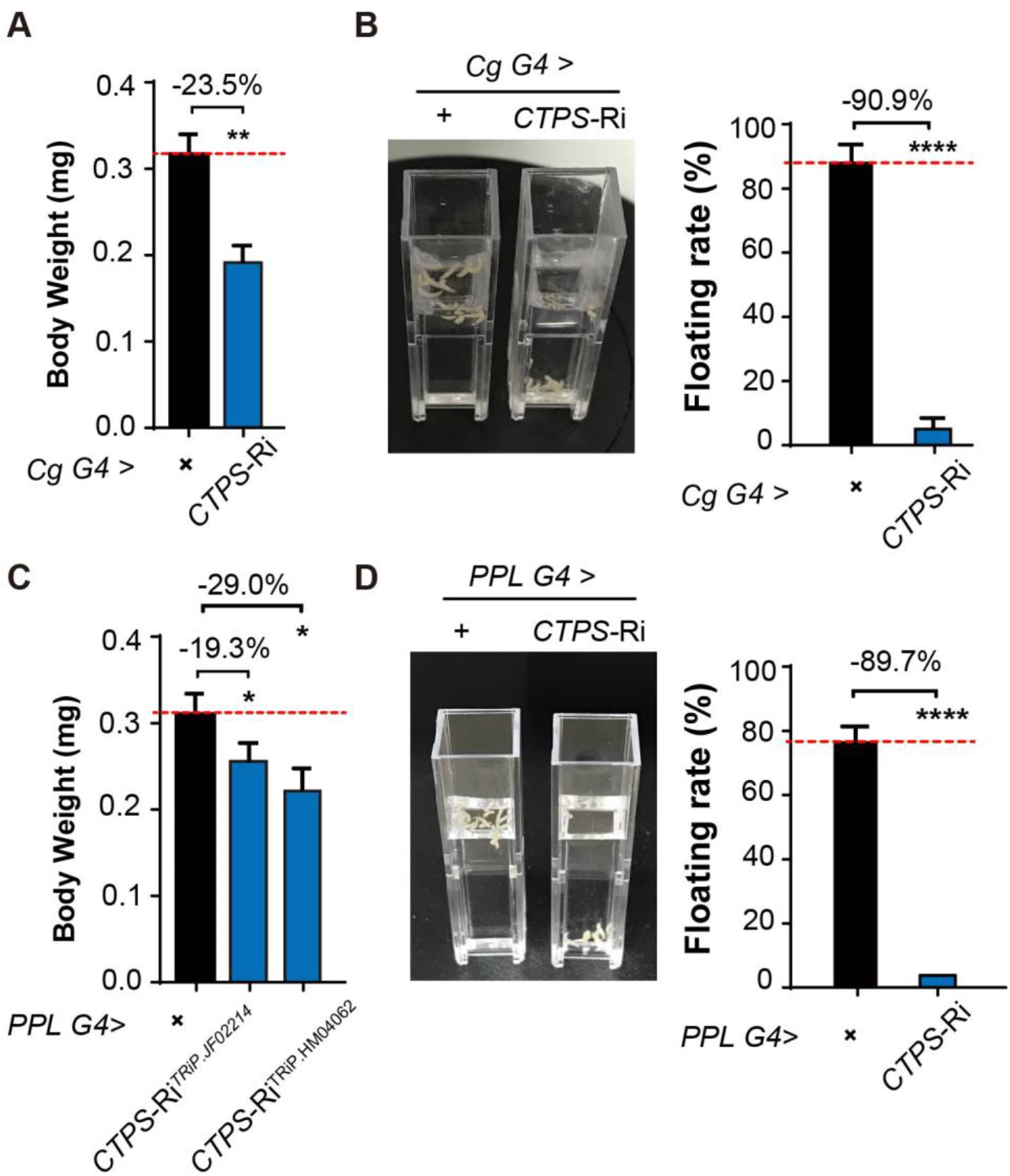
Adipocyte-specific knockdown of *CTPS* decreases larval body weight. (**A**) The early third instar larvae body weight of the indicated lines is measured. (30 larvae/group, 5-6 groups/genotypes). *Cg G4>CTPS*-Ri versus *Cg G4>+*. (**B**) The representative image of floating assay. The quantification of floatation scores (% floating larvae, right panel). *Cg G4>CTPS*-Ri versus *Cg G4>*+ (10 larvae/group, 5-6 groups/genotypes, 3 biological replicates). (**C**) The early third instar larvae body weight of the indicated lines was measured. (30 larvae/group, 5-6 groups/genotypes, 3 biological replicates). *PPL G4>CTPS-Ri* versus *PPL G4>* +. (**D**) The representative image of floating assay. The quantification of floatation scores (% floating larvae, right panel). *PPL G4>CTPS-Ri* versus *PPL G4>* +. (10 larvae/group, 5 groups/genotypes, 3 biological replicates). Data are shown as means ±S.E.M. * P<0.05, **p < 0.01, ***p < 0.001, ****p < 0.001 by Student’s t test compared to the control.

To rule out potential off target, we also used another *CTPS* RNAi line, *CTPS-* Ri^TRip^.^JF02214^. Both body weight (Figure 2C) and larvae floating rate (Figure 2D) in *PPL G4*>*CTPS*-Ri^TRiP.HM04062^ or *PPL G4>CTPS-Ri^TRIP.02214^* were dramatically reduced in either *PPL G4*>*CTPS*-Ri^TRiP.HM04062^ or *PPL G4* >*CTPS*-Ri^TRiP.JF02214^ compared with *PPL G4>+* lines. In addition, we did not found any visible developmental delay in all larval developmental stages in *Cg G4>CTPS-Ri*.

### HFD promotes CTPS expression in the fat body

Given fat body CTPS’s ability to regulate body weight in both adults and larvae, we asked what changes of CTPS cytoophidia occur in adipocytes response to metabolic adaptation to HFD feeding. We challenged mCherry and V5 tagged *CTPS* knock-in larvae (*CTPS-mCh*) with high fat food feeding (HFD, 30% coconut oil) to increases lipogenesis in fat body. We firstly examined the expression level of *CTPS* in *w^1118^* larval flies upon HFD feeding. Quantitative RT-PCR analysis revealed that *CTPS* in the fat body is up-regulated by 120% under HFD feeding compared to regular food (RD) condition (Figure 3A). Moreover, CTPS cytoophidia was elongated up to 60% (Figure 3B, C, D) in *CTPS-mCh* larval fat body, as well as slight increase in numbers of cytoophidia under HFD condition compared to that of RD feeding (Figure 3E).

**Figure 3.**
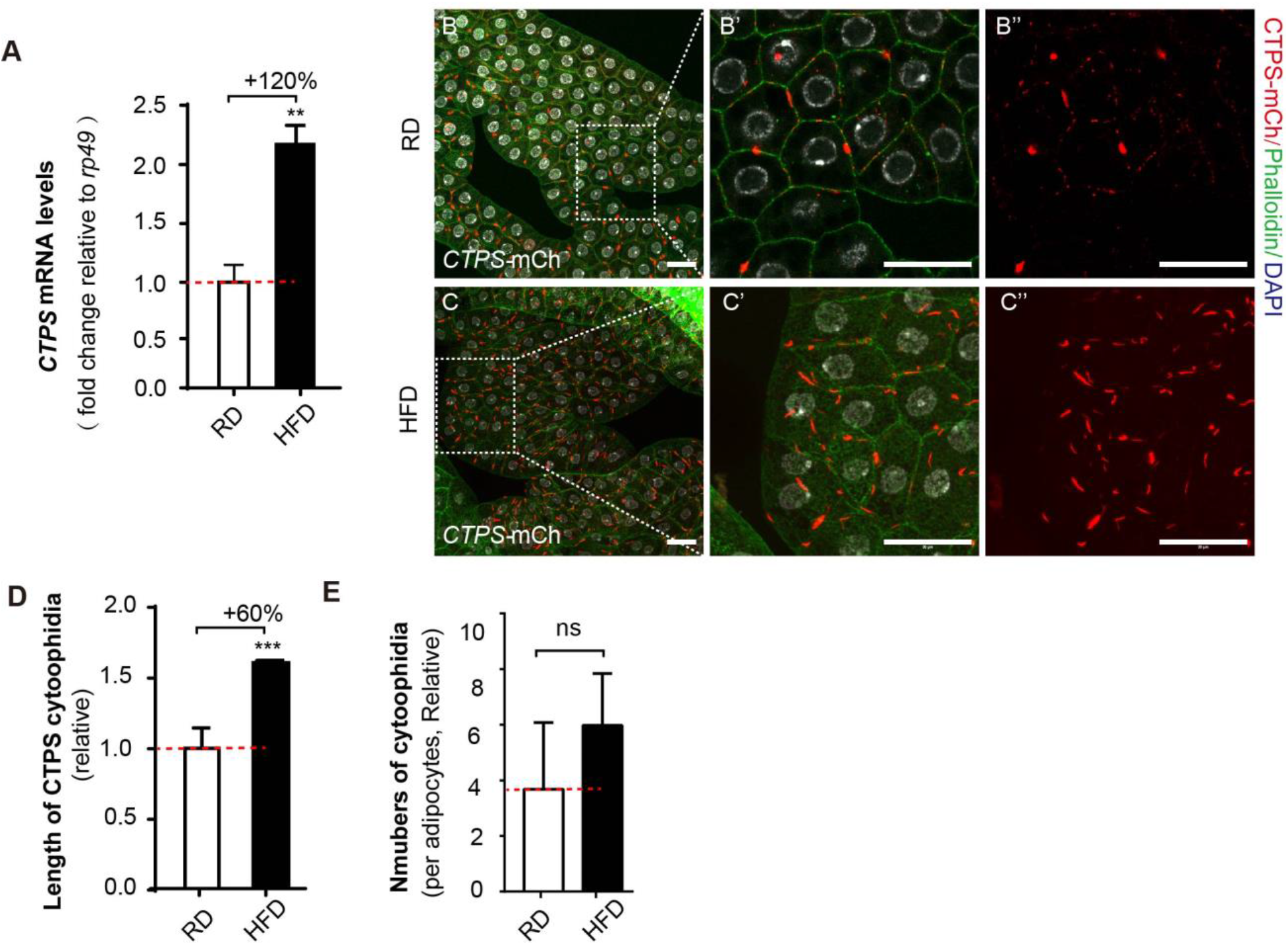
HFD promotes CTPS expression in the fat body. (**A**) Quantitative RT-PCR analysis of the mRNA abundance of *CTPS* from the fat body lysates of early 3^rd^ instar larvae under RD and HFD condition. The relative value is normalized with larvae under RD feeding. (**B**-**C**) Representative confocal images of fat bodies from the early 3^rd^ instar larvae show that CTPS cytoophidia elongated upon HFD feeding (C, C’, and C”) compared to RD condition (B, B’, and B’’). The area marked by the white square is magnified in the right panel (B’, B’’, C’, and C’’). The plasma membrane of the cells is stained with phalloidin (Green). Nuclei are stained with DAPI (Gray). Scale bar 20μm. (**D**) Quantification of the length of cytoophidia shown in B-C. The relative value is normalized with larvae under RD feeding (20 images/genotypes, 3 biological replicates) (**E**) Quantification of the numbers of cytoophidia per adipocyte is shown in B-C. The relative value is normalized with larvae under RD feeding (20 images/genotypes, 3 biological replicates). All values are the means ±S.E.M. ns, no significance, **\*\*** p < 0.01, **\*\*\*** p < 0.001, by Student’s *t*-test.

### Fat body-specific knowdown of CTPS alleviates HFD-induced obesity

Next we investigated whether CTPS acts through the fat body to regulate metabolic adaptation and lipogenesis. We conducted loss of function study by interfering CTPS with RNAi in larval adipose tissue, by *Cg G4*. Response to HFD feeding, the GFP fluorescence intensity of wild-type larvae exhibited an increase reflecting an increase in fat storage suggesting that HFD-induced elevate lipogenesis (Figure 4A). Importantly, fluorescence intensity of *Cg G4>CTPS-Ri* larvae did not exhibit an obvious increase upon HFD feeding compared to that of larvae under RD condition (Figure 4A). Subsequently, upon HFD feeding, *Cg G4>+* larvae exhibited significant body weight gain (11.1%, 0.338 mg; S.E.M.: ± 0.023 mg) compared to that (0.304 mg; S.E.M.: ± 0.018 mg) of RD condition (Figure 4B). Notably, compared to HFD-fed *Cg G4>+* flies, HFD-fed *Cg G4>CTPS-Ri* flies (0.234 mg; S.E.M.: ± 0.018 mg) showed 30.7% reduction of body weight and 23.4% lower body weight gain (Figure 4B).

**Figure 4.**
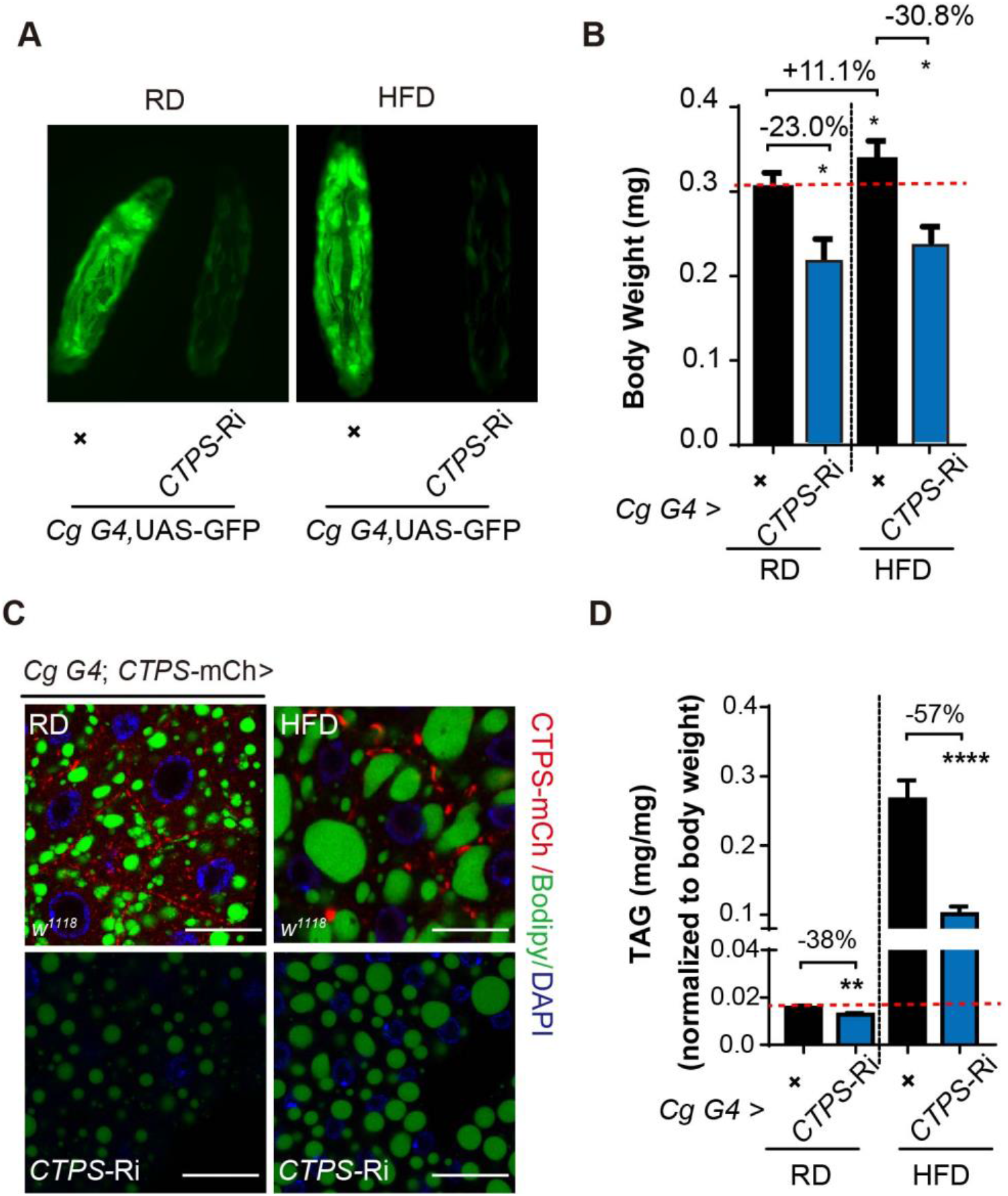
Knockdown of *CTPS* in adipocytes alleviates HFD-induced obesity. (**A**) Compared with the wild-type control, the early 3^rd^ instar larvae expressing GFP (green) with *Cg G4* driving *CTPS* knockdown in the fat body were fed with RD or HFD. (**B**) The body weight of the early 3^rd^ instar larvae in *Cg G4>CTPS-Ri* and *Cg G4>+* is measured under the condition of RD and HFD. (10-30 larvae/group; 5-6 groups/genotypes, 3 biological replicates). (**C**) The lipid droplets of the early 3^rd^ instar larvae are analyzed by confocal microscopy under RD and HFD. Lipid droplets are stained with BODIPY 493/503 and nuclei were stained with DAPI. Scale bars, 20 μm. (**D**) The triacylglycerol (TAG) level of the early 3^rd^ instar larvae in *Cg G4>CTPS-Ri* and *Cg G4>+* under RD and HFD conditions. TAG is normalized to body weight (10 larvae/group; 5-6 groups/genotype, 3 biological replicates). Data are shown as mean ± S.E.M. * P < 0.05, ** P < 0.01, **** P < 0.0001, by one-way ANOVA with a Tukey *post hoc* test.

Furthermore, we imaged lipid droplets with BODIPY 493/503 staining and examined the TAG contents of larvae. Upon HFD feeding, *Cg G4>+* flies exhibited enlarged lipid droplets in the fat body. Notably, compared with the control group, the *Cg G4>CTPS-Ri* showed much fewer and smaller lipid droplets in adipocytes under both RD and HFD conditions (Figure 4C). Indeed, TAG content was decreased by 38% under RD condition, and 57% under HFD condition, in *Cg G4>CTPS-Ri* compared with *Cg G4>+* control group (Figure 4D). These results suggest that CTPS deficiency causes reduction in lipogenesis which may lead to body weight loss of larvae.

### Fat body-specific knockdown of CTPS reduces lipogenic gene expression

We next explored putative CTPS-dependent molecular pathways impacting adipocyte function and lipid homeostasis. First, we performed genome wide RNA sequencing (RNA-Seq) of wild type (*Cg G4*>+) and CTPS deficiency (*Cg G4*>*CTPS*-Ri) larval adipocytes. This revealed significant differences in RNA transcriptional profile with 273 genes. Of them, 204 genes were up-regulated (74.7%) and 69 genes (25.3%) were down-regulated compared to the control (≥2.0 fold change, Student’s t-test, P<0.05) (Figure 5A-C).

**Figure 5.**
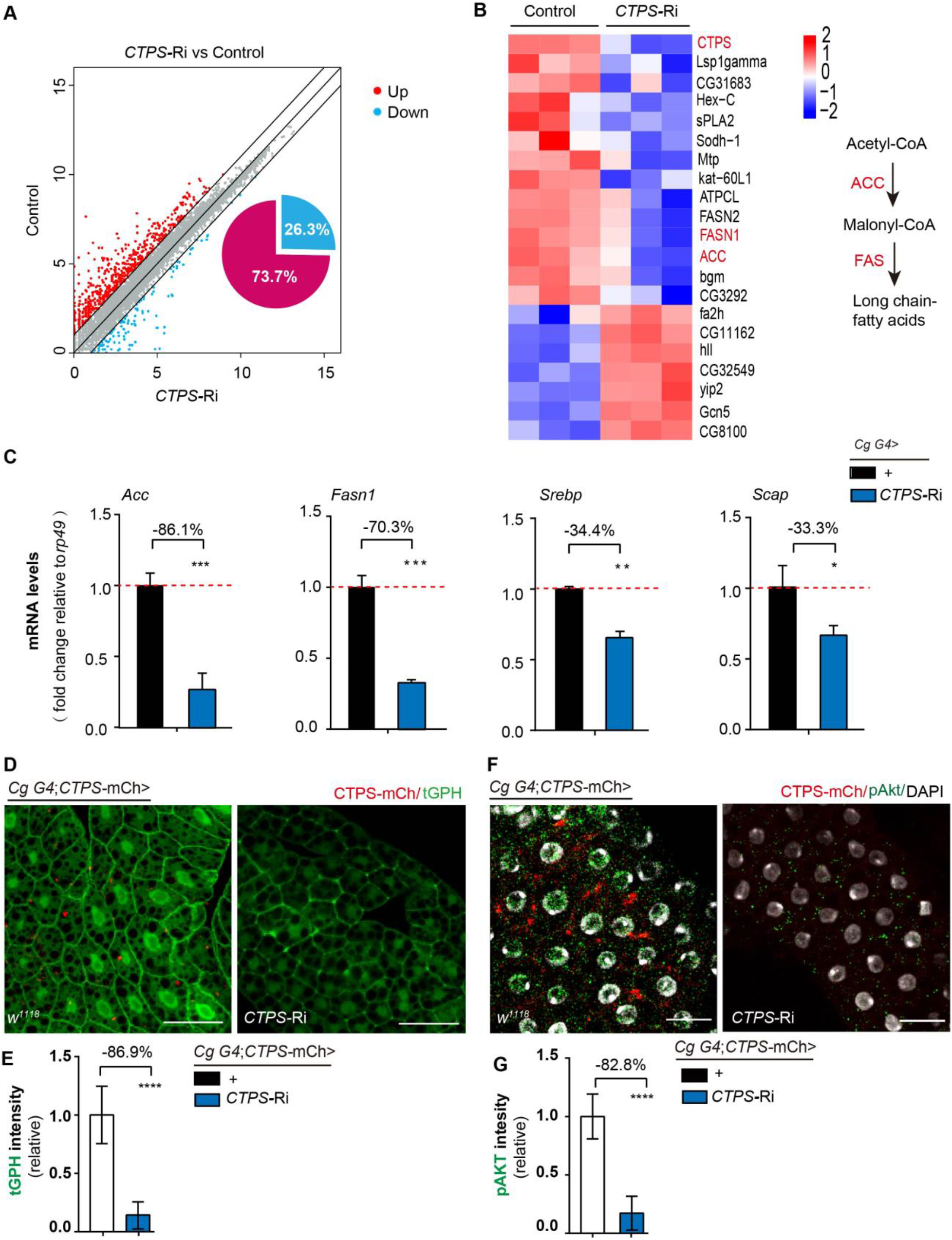
Fat body-specific knockdown of CTPS reduces lipogenic gene expression. (**A**-**B**) Fat body of early 3^rd^ instar larvae feeding under RD condition from *Cg G4>CTPS-Ri* and *Cg G4>+* is analyzed by RNA-seq analysis. A scatter plot showing comparison of transcript abundance between knockdown *CTPS* and the control. On the basis of the comparison of fold change values (gene expression level in the *Cg G4>CTPS-Ri* relative to the *Cg G4>+* control line, ≥ 2.0 fold change, Student’s t-test, P< 0.05). Each dot represents one protein-coding gene (A). A heat map of relative gene expression is depicted for transcripts encoding central enzymes in lipid metabolism from controls (left) and *CTPS* knockdown (right) using RNA-seq data (B). The schematic of long chain-fatty acids synthesis pathway on the right panel. (**C**) Quantitative RT-PCR analysis of the mRNA abundance of *Acc, Fasn1, Srebp* and *Scap* from the fat body lysates of the early 3^rd^ instar larvae in *Cg G4*>*CTPS*-Ri and *Cg G4*>+ flies (30 larvae/group; 5-6 groups/genotype, 3 biological replicates). (**D**) Representative confocal images of PI3K activation in the fat bodies of the 3^rd^ instar larvae. The location of tGPH shows the activity of PI3K. Scale bars, 50 μm. (**E**) The ratio of GFP intensity of the membrane to plasma from D. The value is normalized to the control (10 images/genotypes, 3 biological replicates). (**F**) Representative immunofluorescent staining of phosphorylated Akt in early 3^rd^ instar larvae fat body cells. Phosphorylated Akt are stained with p-Akt (Thr308) antibody. Nuclei are stained with DAPI. Scale bars, 10 μm. (**G**) The GFP intensity of phosphorylated Akt from F. The value is normalized to the control (10 images/genotypes, 3 biological replicates). All data are shown as mean ± S.E.M. * P < 0.05, ** P < 0.01, *** P < 0.001, **** P < 0.0001 by Student’s *t*-test.

Kyoto Encyclopedia of Genes and Genomes (KEGG) analysis revealed that the differently regulated genes by *CTPS* were significantly enriched in genes encoding proteins that were involved in lipid metabolism, carbohydrate metabolism, amino acid metabolism (Figure S2). Of particular interest was the down-regulated expression of lipogenic enzyme genes encoding Acetyl-CoA carboxylase (ACC) and fatty acid synthase 1 (FASN1) in *CTPS* knockdown larvae fat body, and several other genes encoding proteins involved in lipid metabolism (Figure 5C).

To validate these data, we detected the expression level of genes by quantitative RT-PCR. We found that *CTPS* knockdown reduced the expression levels of *Acc* and *Fasn1* by 42%, and 70.3%, respectively (Figure 5C).

One important regulator of lipogenesis is sterol regulatory element binding proteins (SREBP), a highly conserved, membrane bound, transcription factor [26]. In the ER, active SREBP forms complex with SREBP cleavage-activating protein (SCAP), which translocate into the nucleus and promote transcription of genes involved in lipid metabolism such as *Acc* and *Fasn1*. Then we asked whether CTPS knock down also inhibited SREBP transcriptional activity. We tested this by RT-PCR. Expression levels of *Srebp* and *Scap* were reduced by 34.4% and 33.3%, respectively (Figure 5C).

### Fat body-specific knockdown of CTPS suppresses PI3K activity

Phosphatidylinositol 3’-kinase (PI3K) is involved in the phosphorylation of membrane inositol lipids, thus mediating intracellular signal transduction. Activation of PI3K can promote the growth and metabolic activity of fat body [27]. tGPH was used as a cytological marker to monitor the PI3K activity of larval adipocytes [28]. We found that tGPH signal of the cytosol to the plasma membrane is decreased in the early 3^rd^ instar larval fat body of *Cg G4*>*CTPS*-Ri (Fig 5D, E).

Akt is a direct target of PI3K signal. We hypothesized that with the reduction of plasma membrane PI3K activity, *CTPS* deficiency may impair phosphorylation of Akt. We tested this by immunostaining Akt. The Akt signals exhibit punctate patterns in both the cytosol and cell periphery in wild type adipocytes. Notably, the Akt signals are abolished in *CTPS* knockdown adipocytes, indicating a decreased membrane recruitment of Akt in *Cg G4>CTPS*-Ri (Figure 5F,G).

### Disrupting filament-forming property of CTPS alleviates HFD-induced obesity

Then we asked what function CTPS cytoophidia might serve in adipocytes in response to HFD feeding. We identified that H355 of CTPS in the domain of glutamine amidotransferase (GAT) is critical for *Drosophila* cytoophidium formations[29, 30]. We generated a transgenic fly lines which harbored mCherry-HA tag wild type CTPS or H355A point mutation CTPS. Using the *Cg G4* driver, we specifically overexpressed wild-type (CTPS^WT^) or mutant CTPS (CTPS^H355A^) in adipocytes.

Overexpression of the H355A mutant CTPS dominantly disrupted cytoophidia formation in adipocytes (Figure 6D). Moreover, overexpression of the H355A mutant CTPS inhibited HFD-induced increases in body weight (Figure 6A, B) and TAG content (Figure 6C) compared with the wild-type control. The quantification showed that 41.4% body weight gain in *Cg G4*>CTPS^WT^-OE (0.478 mg; S.E.M.: ± 0.026 mg) compared to wild type control (0.338 mg; S.E.M.: ± 0.023 mg) under HFD feeding (Figure 6B). In contrast, 33.7% body weight loss in *Cg G4*>CTPS^H355A^-OE (0.224 mg; S.E.M.: ±0.018 mg) in compared to wild type control under HFD feeding (Fig. 5B). Subsequently, TAG content was reduced by 36.8% in *Cg G4* >CTPS^H355A^-OE (Figure 6C).

**Figure 6.**
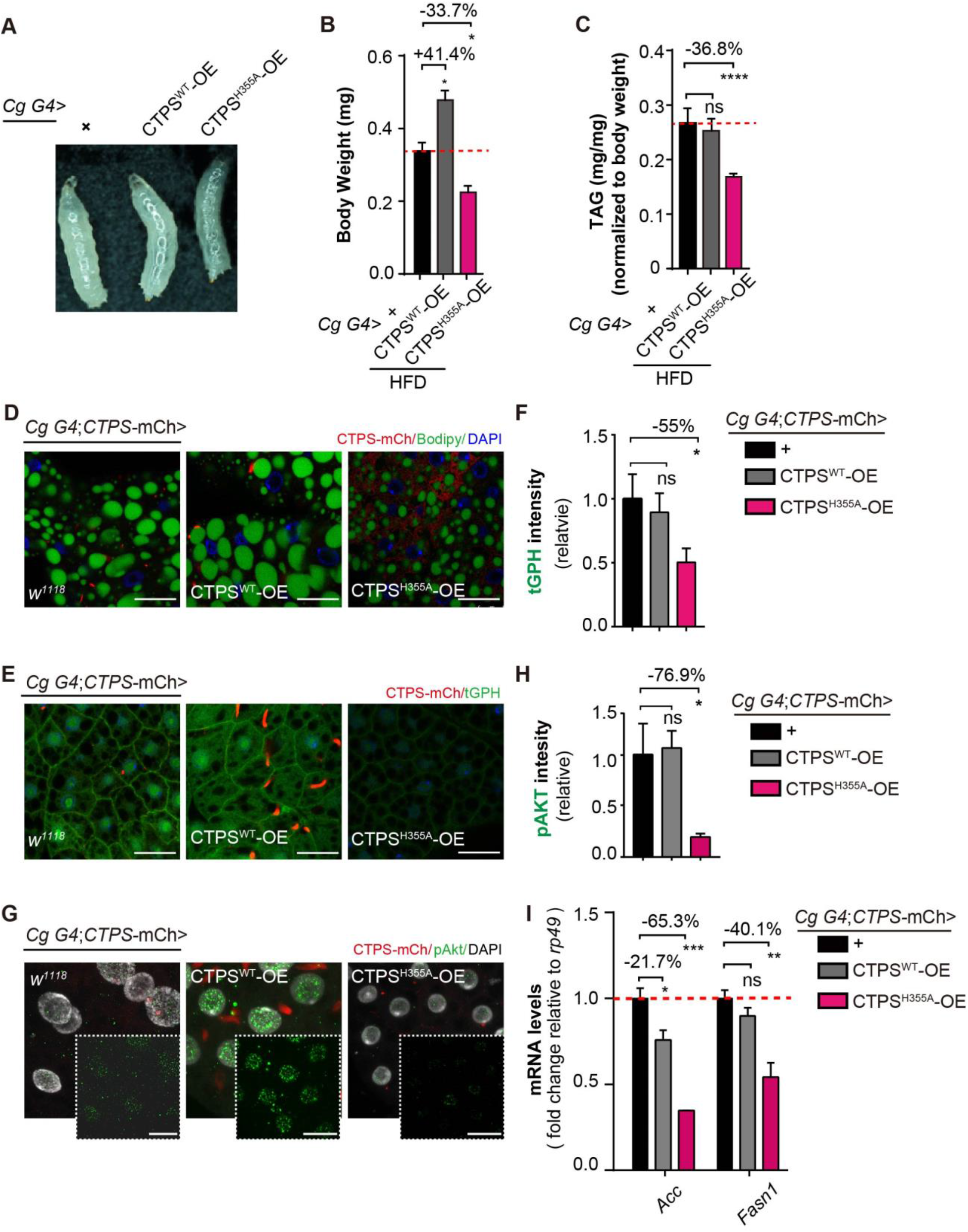
Disrupting filament-forming property of CTPS alleviates HFD-induced obesity. (**A**) Representative images of the early 3^rd^ instar larvae showing larvae morphology where overexpressing mutation H335A (right) compared to that overexpressing wild type CTPS (middle) and control (left) by *Cg G4* driver under HFD condition. (**B**) The body weight of the third instar larvae is measured under HFD condition. (30 larvae/group; 5-6 groups/genotypes, 3 biological replicates). (**C**) Under HFD condition, the TAG level of the early 3^rd^ instar larvae of *Cg G4*>CTPS^WT^-OE, *Cg G4*>CTPS^H355A^-OE and *Cg G4*>+ line. TAG is normalized to body weight (10 larvae/group; 5-6 groups/genotypes, 2 biological replicates). (**D**) Representative confocal images of lipid droplets in the fat bodies of the 3^rd^ instar larvae (*Cg G4*>CTPS^WT^-OE, *Cg G4*>CTPS^H355A^-OE and *Cg G4*>+) under RD or HFD condition. Scale bars, 20 μm. (**E**) Representative confocal images of PI3K activation in the fat bodies of the 3^rd^ instar larvae. The location of tGPH shows the activity of PI3K. Scale bars, 20 μm. (**F**) The ratio of GFP intensity of the membrane to cytosol from E. The value is normalized to the control (10-15 images/genotypes, 3 biological replicates). (**G**) Representative immunofluorescent staining of phosphorylated Akt in early 3^rd^ instar larvae fat body cells. Scale bars, 20 μm. The phosphorylated Akt staining (green) channels were shown in white boxes. (**H**) The GFP intensity of phosphorylated Akt from G. The value is normalized to the control (10 images/genotypes, 3 biological replicates). (**I**) Quantitative RT-PCR analysis of *acc and fasn1* mRNA abundance in fat body lysate of the 3^rd^ instar larvae from the indicated genotypes (30 larvae/genotype; 4 groups/genotypes, 2 biological replicates). All data are shown as mean± S.E.M. ns, no significance, * P < 0.05, ** P < 0.01, *** P < 0.001, **** P < 0.0001 by Student’s *t*-test or one-way ANOVA with a Tukey *post hoc* test.

Confocal images of BODIPY 493/503 staining revealed that both *Cg G4; CTPS*-mCh>CTPS^WT^-OE and the control group increased lipid droplets accumulation, whereas much fewer and smaller lipid droplets in the fat of *Cg G4; CTPS*-mCh>CTPS^H355A^-OE flies in the response to HFD (Figure 6D).

We found that the fat body containing H355A mutant resulted in a significant decrease of tGPH-associated fluorescence on the adipocyte membrane (Figure 6E), there was 50.3% reduction of tGPH signal of the cytosol to the plasma membrane (Figure 6F), indicating a dramatic down-regulation of PI3K activity in the absence of cytoophidia. Furthermore, the Akt signals was diminished in *Cg G4*>CTPS^H355A^*-*OE adipocytes (Figure 6G, H). Of note, we found that compared with *Cg-G4*>+, RT-PCR showed a 65.3% and 40.1% decrease in *Acc* and *Fasn1* expression level in *Cg G4*>CTPS^H355A^-OE, while no obvious changes of *Fans1* mRNA levels in *Cg G4*>CTPS^WT^-OE in respect to the control (Figure 6I).

## Discussion

Obesity has become an epidemic disease globally, with more than 1.9 billion adults overweight and 650 million of them considered obese in 2016, 38 million children (under the age of 5) overweight or obese in 2019. *Drosophila* is emerged as an ideal model for lipid metabolism and homeostatic regulation [18, 31], with evolutionarily and functionally conserved metabolic signaling pathway and CTPS cytoophidia [24]. We identify that CTPS cytoophidia in the fat body links to lipid homeostasis and HFD-induced obesity. Our results demonstrated that CTPS cytoophidia elongated in response to HFD feeding and affected diet-induced lipid accumulation.

CTPS catalyzes the rate-limiting step which converts UTP to CTP. CTPS filament formation was most frequently observed, when cells went into stationary phase [7] or were in depletion of Glutamine in cancer cells [32], which promotes cancer cell growth after stress alleviation. This suggests a functional association with the energy-fluctuated cellular state, an assumption that has so far remained untested.

*Drosophila* fat body is a crucial organ to regulate lipid deposit in response to HFD feeding. In this study, we found that CTPS deficiency specifically in the fat body decreased body weight and sensitized flies to starvation during food deprivation, indicating that CTPS may act mainly in the fat body to regulate systemic energy homeostasis. Importantly, we found that HFD feeding leaded to augment CTPS transcription and elongated cytoophidia. This suggests that CTPS cytoophidia responses to changes of nutrient availability. It has documented that polymerization of CTPS regulates enzymatic activity [33, 34] and modulates protein stability [16]. Dynamic regulation of CTPS assembly may explain why CTPS cytoophidia elongate in adipocytes upon HFD feed.

Lipid metabolism within *Drosophila* adipocytes is tightly regulated. Here, we find that loss of CTPS by RNAi interference or ectopic expression of H355A mutant CTPS suppresses PI3K activity and reduces phosphorylated Akt level. Akt, as a downstream effector of PI3K, appears to stimulate *de novo* lipid synthesis through the activation of SREBP [36–38]. In *Drosophila*, only single SREBP homolog, a center regulator of lipid metabolism, targets lipogenic genes such as *Acc*, *Fasn* [26]. *Fasn* is the key metabolic multienzyme that is responsible for the terminal catalytic step in fatty acid synthesis. The genetic variation within the Human *FASN* is associated with obesity in adults [39]. High transcriptional activation of *FASN* are also documented in the cancer cells [40–42]. In this study, either CTPS depletion or overexpression of H355A mutant CTPS results in down-regulated lipogenic enzymes (*Acc*, *Fasn*), which might be due to suppressing PI3K-Akt signaling.

Similarly, FASN protein expression can also be inhibited by pharmacological blockade of PI3K signaling in cancer cells [43]. Given similarity of *Drosophila* fat body and mammalian liver and adipose tissue, it would be interesting to investigate whether CTPS deficiency-mediate suppression of lipid storage via the PI3K/Akt-FASN axis may operate in mammalian adipose tissues and other tissues.

## Materials and Methods

### Generation of transgenic flies

The CRISPR/Cas9 technology was used to establish the C-terminal mChe-4V5 tagged CTPS knock-in fly according to homology-directed repair procedures as previously described [44] at Fungene Biotech (http://www.fgbiotech.com). The two of single strain guide RNA (sgRNA) were designed with CRISPR Optimal Target Finder [45]. sgRNA1: GCCATAAGTAAACTAGTGAACGG, sgRNA2: CCTAAAGTGTTTAACATCCGATT. For donor vector construction, the mChe-4V5 cassette amplified from pBluescript SK-mCherry-4V5 (from Fungene Biotech, unpublished) was cloned into pBSK (-) vector containing CTPS arm regions. The donor vector pBSK CTPS-mCherry-4V5, sgRNAs and cas 9 mRNA were injected into *w^1118^* embryos. PCR sequencing was performed to validate if the offspring flies carried mCherry-4V5 insertion.

To generate transgenic UAS-CTPS and UAS-CTPS^H355A^ flies, the cDNAs encoding *Drosophila CTPS* was produced by RT-PCR using the total RNAs extracted from the *w^1118^* line (#3605, from the Bloomington *Drosophila* Stock Center). The oligonucleotide primers used were as follows: *CTPS* sense 5’-ttcgttaacagatctgcggccgcatgaaatacatcctggtaact-3’, antisense 5’-ttcacaaagatcctctagaggtacccttgtacagctcgtccatgc-3’. The PCR products were digested with NotI and KpnI and cloned into the pUASTattB vector for the expression of mCherry-HA-tagged CTPS protein. H355A point mutation was introduced into pUAST-attB-CTPS plasmid by using Mut Express II Fast Mutagenesis Kit V2 (Vazyme) following the manufacturer’s instructions. The mutagenic oligonucleotide primers used was sense 5’GAGCAAGTACCGGAAGGAGTGGCAGAAGCTATGCGATAGCCAT-3’, antisense 5’-TGCCACTCCTTCCGGTACTTGCTCGGCTCAGAATGCAAAGTTT-3’. Then, site-specific integration of pUASTattB-CTPS or pUASTattB-CTPS^H355A^ plasmid into fly germ line (attp2) that contain attp lading sites was carried out by coinjection with phiC31integrase RNA as previously described [46] at the Core Facility of *Drosophila* Resource and Technology, SIBCB, CAS.

### Fly strains

The G4/UAS system [47] was utilized for adipocyte-specific expression or RNAi knockdown of the desired genes.. The fly lines were obtained from the Bloomington *Drosophila* Stock Center (BDSC; Department of Biology, Indiana University, Bloomington, IN), including *w^1118^* (stock number 3605), *Cg G4* (stock number 7011), *PPL G4* (stock number 5092), *CTPS*-RNAi^TRiPJF02214^ (stock number 31924), *CTPS*-RNAi^TRiP.HM04062^ (stock number 31752), and tGPH (stock number 8163).

### Fly husbandry and diet preparation

Fly lines were raised on standard yeast-cornmeal-agar food. To make a high fat diet, we added 30% coconut oil (v/v) to the regular diet and well mixed. For rearing the flies, 5 mL of the diet was prepared per vial. To make sure larvae used in this study at the desired developmental stage, we restricted egg collections by allowing female to lay eggs for less 4 hours. All flies were cultured at 25°C with 50% humidity under a 12 h/12 h light/dark cycle.

### Fly body weight

The embryos were collected for 2-4 hours and raised on experimental diets. After 72-80 hrs, the larvae were rinsed in PBS and then the larval body weight (30 flies per group, 3-6 groups per genotypes) was measured. For body weight of adults, five-day-old adults were measured (30 flies per group, 3-6 groups per genotypes).

### Starvation assay

For starving the adult fly, adults at 5-day-old were placed in vial with 3 ml 1% agar (about 30 flies per vial, 5-6 groups per genotype) and were transferred every two days to avoid bacterial contamination. Dead flies were quantified every 12 hours as the time until death.

### Floating assay

Floating assays were performed as previously described [20, 48]. Briefly, 10 3^rd^ instar larvae in each experimental group were put in 10 ml of 9% sucrose, PBS solution. Larvae were placed in the sucrose solution, gently mixed and, at a 3 minutes time point, the larvae floating at the surface of solution were counted. The data represents the percentage of floating larvae. The experiments were repeated at least three times.

### Triglyceride analysis

The 3^rd^ instar larvae were snap frozen in liquid nitrogen and stored at −80°C. Each biological replicate represents 10 flies collected into a 1.5-ml microcentrifuge tube, and each experiment includes six biological replicates for each genotype. Triglyceride concentration was measured using a coupled colorimetric assay as previously described [20]. Samples were homogenized in phosphate-buffered saline (PBS) with 1% Triton-X and immediately incubated at 70 °C for 10 min. Heat-treated homogenates were incubated with Free Glycerol Reagent (Sigma) for 5 min at 37 °C. Samples were assayed using microplate spectrophotometer at 540 nm. TAG levels were normalized to body weight of larvae.

### Immunohistochemistry

The fat body from early 3^rd^ instar larvae were dissected in Grace’s Medium and then fixed in 4% formaldehyde in PBS for 15 min before immunofluorescence staining. For membrane staining, fixed fat bodies were washed twice for 5 min in PBS and then were incubated with 0.165 μM Alexa Fluor 488 phalloidin (Invitrogen) in PBSTG for 30 min at RT. Then samples were rinsed in PBS twice for 5min each and mounted in Vecta shield with DAPI (Invitrogen). For lipid droplet staining, fixed fat bodies were washed twice for 5 min in PBS and then were incubated with BODIPY 493/503 (1 μg/mL for 30min at RT). Then samples were rinsed in PBS twice for 5min each and mounted in Vecta shield with DAPI (Vector Labs). The primary antibody used is rabbit anti-Phospho-Akt (Thr308) (244F9) antibody (1:500, Cell Signaling, Catalogue no. 4056). Secondary antibody is anti-rabbit IgG, (1:2000, Cell Signaling, Catalogue no.5151).

### Imaging and image analysis

Fluorescent images were obtained by confocal laser-scanning microscopy (Leica SP8). Super resolution images were obtained by TCS SP8 STED 3X super resolution stimulated emission depletion microscopy. For quantification of length of CTPS-mCherry cytoophidia, the length of CTPS cytoophidia was counted in cells of 40X confocal images with FIJI-ImageJ. For quantification of numbers of CTPS-mCherry cytoophidia, the number of CTPS mCherry-containing cytoophidia was counted in cells of 40X confocal images with FIJI-ImageJ. The data represents cytoophidia number in one cell normalized by cell number in one such image.

### RNA isolation and RNA sequencing

Briefly, total RNA was isolated from 50–100 of 2^nd^ instar larvae of dissected fat body, using TRIzol Reagent (Invitrogen) according to the manufacturer’s recommendations. The libraries were constructed using TruSeq Stranded mRNA LT Sample Prep Kit (Illumina, San Diego, CA, USA) according to the manufacturer’s instructions. P value < 0.05 and fold Change > 2 or fold Change <0.5 was set as the threshold for significantly differential expression. The transcriptome sequencing and analysis were conducted by OE Biotech Co., Ltd. (Shanghai, China).

### Quantitative RT-PCR

Total RNAs were prepared from second-instar larvae from using the TRIzol reagent (TransGen Biotech, Beijing, China). cDNAs were synthesized with PrimeScript RT Master mix (Takara) followed by adding template RNA. 2X SYBR Green PCR Master Mix was purchased from Bimake. Real-time quantitative PCR was conducted using the QuantStudion^TM^ 7 flex System (Applied Biosytstems). For normalization, *rp49* was utilized as the internal control. The oligonucleotide primers used were as follows: *rp49:* sense 5’-TCCTACCAGCTTCAAGATGACC-3’, antisense 5’-CACGTTGTGCACCAGGAACT-3’; *CTPS:* sense 5’-GAGTGATTGCCTCCTCGTTC-3’, antisense 5’-TCCAAAAACCGTTCATAGTT-3’. *Acc*: sense 5’-GTGCAACTGTTGGCAGATCAGTA-3’ antisense 5’-TTTCTGATGACGACGCTGGAT-3’ *Fasn1*: sense 5’-CCCCAGGAGGTGAACTCTATCA-3’ antisense 5’-TTTCTGATGACGACGCTGGAT-3’ *Srebp:* sense 5’-GGCAGTTTGTCGCCTGATG-3’ antisense 5’-CAGACTCCTGTCCAAGAGCTGTT-3’ *Scap:* sense 5’-ACCAGAGCAGCGAAAACAAAC-3’ antisense 5’-GAGAGTTCTGCGTCCACAGG-3’

### Statistical analysis

All data are presented as the mean ± standard errors of the mean (S.E.M) from at least three independent experiments. Statistical analysis between each genotype and the controls was determined by unpaired two-tailed Student’s *t*-test, whereas multiple comparisons between genotypes were determined by one-way ANOVA with a Tukey *post hoc* test in GraphPad Prism 7.0. *P*<0.05 was considered to be statistically significant.

## Acknowledgments

We thank Xiaoming Li from the Molecular Imaging Core Facility (MICF) at School of Life Science and Technology, ShanghaiTech University for providing technical support. We also thank the Core Imaging Facility at the National Center for Protein Science Shanghai (NCPSS).

## Funding

This work was supported by grants from National Natural Science Foundation of China (No. 32071144 and 31771490) to J.L. and J-L.L

## Author contributions

J.L. conceived and designed the studies, analyzed the data. J.L., Y. Zhang, Y. Zhou, and Q-Q.W. performed the experiments. J-L.L. supervised the project. J.L. and J-L.L. wrote the manuscript.

## Competing interests

Authors declare no competing interests.

## Data and materials availability

All data are provided in the text or supplementary materials.

## Supplemental Figures S1, S2 and Figure Legends

**Supplemental Figure S1.**
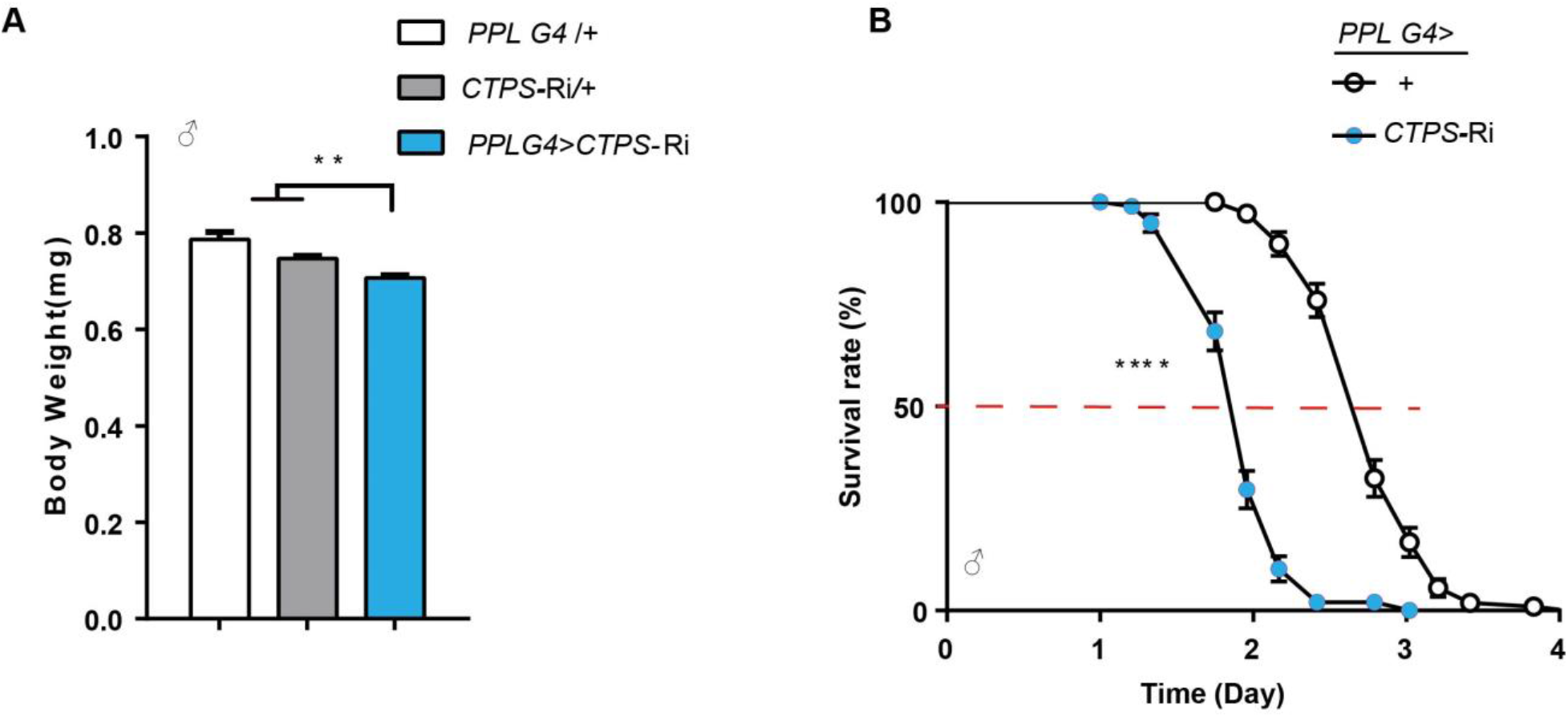
Adipocyte-inhibition *CTPS* decreased body weight and starvation resistance. (**A**) Body weight of 5-days old male adults fly from the indicated genotypes. (30 flies/group, 3 groups/genotype, 2 biological replicates). *PPL G4*>CTPS-Ri versus *PPL G4*>+ or *CTPS-Ri/+*. All values are the means ±S.E.M. **\*\*** p < 0.01, by Student’s *t*-test. (**B**) Survival rates are measured for starved male adult flies from the indicated genotypes (5 days of age; 30 flies/group, 5 groups/genotype, 2 biological replicates). *χ*^2^=162 for male, *PPL G4*, GFP*>* + versus *PPL G4*, GFP*>CTPS-*Ri, **** p < 0.001 by log-rank test

**Supplemental Figure S2.**
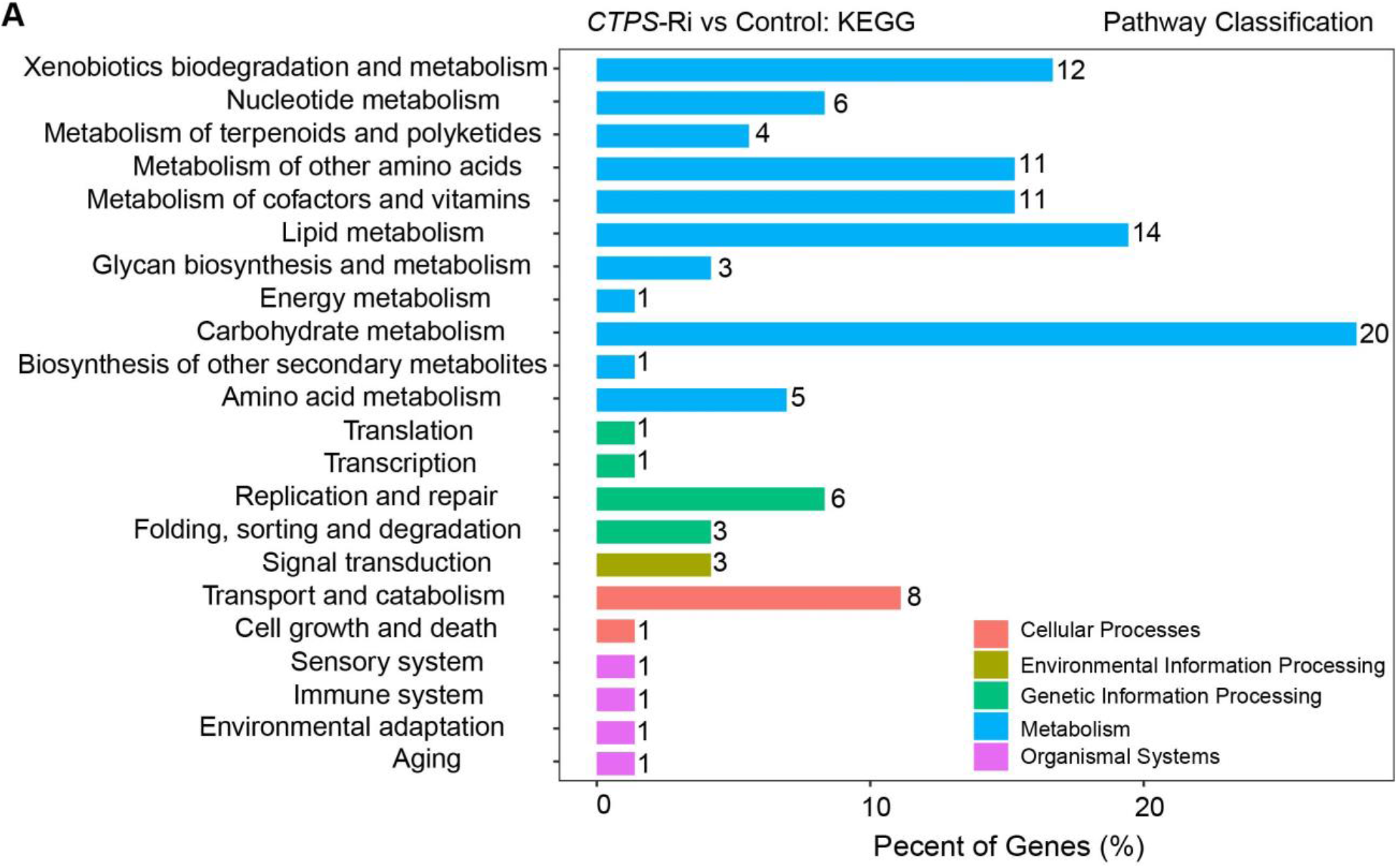
KEGG functional classification of affecting genes upon CTPS knockdown. Numbers: numbers of genes belonging to distinct functional groups which are regulated in the *Cg G4*>CTPS-Ri relative to the *Cg G4*>+ control line. Gene functions are predicted based on KEGG database.

## Notes

### Competing Interest Statement

The authors have declared no competing interest.

